# The contribution of Kaposi’s sarcoma-associated herpesvirus ORF7 and its zinc-finger motif to viral genome cleavage and capsid formation

**DOI:** 10.1101/2022.05.02.490372

**Authors:** Yuki Iwaisako, Tadashi Watanabe, Yuichi Sekine, Youichi Suzuki, Takashi Nakano, Masahiro Fujimuro

**Affiliations:** Department of Cell Biology, Kyoto Pharmaceutical University, Misasagi-Shichono-cho 1, Yamashina-ku, Kyoto, 607-8412, Japan; Department of Microbiology and Infection Control, Faculty of Medicine, Osaka Medical and Pharmaceutical University, Osaka, 569-8686, Japan

## Abstract

Kaposi’s sarcoma-associated herpesvirus (KSHV) is the causative agent of endothelial and B cell malignancies. During KSHV lytic infection, lytic-related proteins are synthesized, viral genomes are replicated as a tandemly repeated form, and subsequently, capsids are assembled. The herpesvirus terminase complex is proposed to package an appropriate genome unit into an immature capsid, by cleavage of terminal repeats (TRs) flanking tandemly linked viral genomes. Although the mechanism of capsid formation in α- and β-herpesviruses are well-studied, in KSHV, it remains largely unknown. It has been proposed that KSHV ORF7 is a terminase subunit, and ORF7 harbors a zinc-finger motif, which is conserved among other herpesviral terminases. However, the biological significance of ORF7 is unknown. We previously reported that KSHV ORF17 is essential for the cleavage of inner scaffold proteins in capsid maturation, and ORF17 knockout (KO) induced capsid formation arrest between the procapsid and B-capsid stages. However, it remains unknown if ORF7-mediated viral DNA cleavage occurs before or after ORF17-mediated scaffold collapse. We analyzed the role of ORF7 during capsid formation using ORF7-KO-, ORF7&17-double-KO (DKO)-, and ORF7-zinc-finger motif mutant-KSHVs. We found that ORF7 acted after ORF17 in the capsid formation process, and ORF7-KO-KSHV produced incomplete capsids harboring non-spherical internal structures, which resembled soccer balls. This soccer ball-like capsid was formed after ORF17-mediated B-capsid formation. Moreover, ORF7-KO- and zinc-finger motif KO-KSHV failed to appropriately cleave the TR on replicated genome and had a defect in virion production. Thus, our data revealed that ORF7 contributes to terminase-mediated viral genome cleavage and capsid formation.

**IMPORTANCE:** In herpesviral capsid formation, the viral terminase complex cleaves the TR sites on newly synthesized tandemly repeating genomes and inserts an appropriate genomic unit into an immature capsid. Herpes simplex virus 1 (HSV-1) UL28 is a subunit of the terminase complex that cleaves the replicated viral genome. However, the physiological importance of the UL28 homolog, KSHV ORF7, remains poorly understood. Here, using several ORF7-deficent KSHVs, we found that ORF7 acted after ORF17-mediated scaffold collapse in the capsid maturation process. Moreover, ORF7 and its zinc-finger motif were essential for both cleavage of TR sites on the KSHV genome and virus production. ORF7-deficient KSHVs produced incomplete capsids that resembled a soccer ball. To our knowledge, this is the first report showing ORF7-KO-induced soccer ball-like capsids production and ORF7 function in the KSHV capsid assembly process. Our findings provide insights into the role of ORF7 in KSHV capsid formation.

## Introduction

KSHV, also known as human herpesvirus 8 (HHV-8), belongs to the Rhadinovirus genus in the *γ-herpesviridae* subfamily (1, 2). KSHV is a major etiological agent of Kaposi’s sarcoma, primary effusion lymphoma, multicentric Castleman’s disease, and KSHV-associated inflammatory cytokine syndrome (2, 3, 4, 5). The tumorigenicity of KSHV has been established, especially within the context of immunosuppressed patients who are undergoing organ transplantation or those who are co-infected with human immunodeficiency virus. KSHV establishes a life-long infection in its host and exists as either a latent or a lytic infection. During latent infection, the KSHV genome circularizes to form an episome in the nucleus. The transition of KSHV between latent infection and lytic replication depends on the immediate early gene ORF50/RTA (replication and transcription activator), which triggers the transcriptional activation of early genes and late genes encoding viral DNA replication enzymes and viral structural proteins (6). The KSHV genome is replicated as a linear form consisting of tandem genomic repeats, and each genomic unit is flanked by a TR sequence (6).

The basic structure and formation of viral capsids are considered to be similar amongst the human herpesviruses because the viral genes related to encapsidation are highly conserved in all human herpesviruses (6). Even though the encapsidation process of HSV-1, which belongs to the α*-herpesviridae* subfamily, is relatively well understood, little is known about the KSHV encapsidation process. In HSV-1 capsid formation, four types of capsids can be detected in the infected cell nucleus during lytic infection. The capsid types are referred to as procapsid, A-capsid, B-capsid, or C-capsid. The characteristic features of each type of capsid are reviewed below (7, 8, 9). Procapsids are constructed on skeletons that contain a large amount of internal scaffold proteins (10). Procapsids are unstable, not angularized, and are spherical rather than icosahedral (11, 12). A-capsids contain little DNA or protein and are considered to be produced by desorption of genome from capsid or unsuccessful genome packaging into a capsid (13, 14). B-capsids comprise a type of angularized capsid morphology as observed by transmission electron microscopy (TEM). B-capsids have a spherical internal structure consisting of scaffold proteins and they lack viral DNA genomes (15). C-capsids, correctly maturated capsids, have an appropriate viral genome unit instead of a spherical internal structure (13, 14). Herpesviral capsid formation occurs via the following steps: (i) procapsid formation, (ii) internal scaffold cleavage by viral proteases, (iii) internal scaffold disruption by detachment from an outer capsid shell, and (iv) mature capsid (C-capsid) formation. Procapsids are formed by the assembly of outer capsid proteins (also referred to as major capsid proteins) and internal scaffold proteins (8). A viral protease cleaves the internal scaffold proteins and the internal scaffold shell is eliminated from the outer capsid shell, resulting in its angularization (16, 17, 18). The HSV-1 terminase complex cleaves the tandemly linked viral genomes and binds to the portal on the capsid, resulting in the packaging of an appropriate genomic unit into an immature capsid (9, 11, 19). Although it is unclear when the terminase-mediated DNA cleavage and DNA packaging into capsids occur, these events are thought to take place between (i) procapsid formation and (iv) C-capsid formation. B-capsids are formed as unsuccessful capsids when the decayed scaffold is not eliminated from the capsid during the aforementioned (i), (ii), and (iii) stages. However, the B-capsid may also be an intermediate capsid in the C-capsid formation process.

In HSV-1, the mechanism of capsid formation has been revealed using recombinant viruses. In contrast, KSHV capsid formation has not been well-studied. We previously showed that the KSHV protease ORF17 was essential for both the cleavage of the internal scaffold protein as well as capsid maturation after procapsid formation (20). The KSHV capsid formation process was arrested between the procapsid and B-capsid stages by viruses lacking ORF17 or viruses lacking ORF17 protease activity (20). Even though procapsids and B-capsids are not generally distinguished by TEM analysis, ORF17-deficient KSHVs produced B-capsid-like procapsids which possessed spherical internal structures as viewed by TEM (20).

HSV-1 terminase is required for viral DNA cleavage and DNA packaging into the immature capsids (9). The terminase cleaves the viral DNA at the TR sites, which flank the tandem repeats of the viral genome. Thus, a correctly cleaved genome is inserted into an immature capsid. Three HSV-1 proteins, UL15, UL28, and UL33 are components of the HSV-1 terminase complex (21, 22). In cells infected with a temperature-sensitive UL28 mutant, capsids containing viral genomes were not observed, but capsids containing compact internal scaffolds were detected (23). Moreover, UL28-deficient HSV-1 is capable of replicating viral DNA, but it is unable to cleave replicated viral concatemer DNA and fails to produce infectious progeny virus (24). Epstein-Barr virus (EBV) BALF3 and KSHV ORF7 are homologs of HSV-1 UL28. EBV BALF3 is required for viral DNA replication and mature capsid formation (25). The putative zinc-finger motif of UL28 (C197-X2-C200-X22-C223-X-H225) is conserved among the UL28 homologs of all herpesviruses, including KSHV ORF7 (25). This motif is essential for HSV-1 terminase function, because recombinant HSV-1 lacking the putative zinc-finger motif of UL28 failed to package viral DNA (21). Thus, the function of terminase and its components has been extensively studied in the context of HSV-1 and EBV. In KSHV, at least three viral proteins (ORF7, ORF29, and ORF67.5) are thought to form the terminase complex because of their protein sequence homology to terminase proteins from other viruses; however, the precise composition of the terminase complex from KSHV remains unknown.

We previously generated an ORF7-KO KSHV bacterial artificial chromosome (BAC) to analyze the role of ORF7 in viral replication. We found that ORF7 was dispensable for viral DNA replication but was essential for virus production (26). Furthermore, ORF7 interacted with ORF29 and ORF67.5, which are homologs of HSV-1 UL15 and UL33, respectively (26). However, it still remains unknown whether ORF7 is a component of the KSHV terminase complex. Specifically, it is unclear whether ORF7 and its zinc-finger motif contribute to viral DNA cleavage. It remains unknown if ORF7-mediated viral DNA cleavage occurs before or after ORF17-mediated scaffold collapse. Therefore, in order to address these questions, we analyzed the virological properties of ORF7 KO-, ORF7&ORF17 DKO-, and ORF7 zinc-finger motif mutant-KSHVs. We found that ORF7 acted after ORF17 in the capsid formation process, and KSHV ORF7 and its zinc-finger motif were required for viral DNA cleavage and the capsid formation.

## Results

### ORF7-KO KSHV produces unique incomplete capsids which resemble soccer balls

We previously generated and analyzed two types of ORF7-KO KSHV, frameshift-induced ORF7 (FS-ΔORF7 or ORF7-fsKO) BAC16 and stop codon-induced ORF7 (ST-ΔORF7 or ORF7-stKO) BAC16. It was revealed that ORF7 is essential for the production of infectious virions (26). However, the role of ORF7 in capsid formation remains unclear. Therefore, to evaluate the contribution of ORF7 in KSHV capsid formation, we observed capsids formed in wild-type (WT) BAC16-, ORF7-fsKO BAC16-, and ORF7-stKO BAC16-harboring iSLK cells (WT-iSLK, ORF7-fsKO-iSLK, and ORF7-stKO-iSLK, respectively) by TEM. The cells were treated with Dox and NaB for 72 h in order to induce a lytic infection via RTA expression. The lytic cells were sampled to analyze the capsids formed in the nucleus (Fig. 1). C-capsids were detected in WT-iSLK (Fig. 1A, indicated by the black arrowhead), and a few A-capsids were also found in the nuclei (Fig. 1A, indicated by the white arrowhead). A small number of B-capsids were also observed in different visual fields (data not shown). Interestingly, many unique capsids which resemble soccer balls were detected in ORF7-fsKO-iSLK and ORF7-stKO-iSLK (Figs. 1B and C, indicated by the white arrows). These incomplete capsids with unique structural features can be considered as a type of B-capsid because they have a non-spherical internal structure within the capsid. The capsids produced in ORF7-KO-iSLK cells contain a common internal structure that appears to be a telstar pattern (coin dot) of the soccer ball. Therefore, we refer to these capsids as ‘soccer ball-like capsids’ in this manuscript. In contrast to WT-iSLK, C-capsids were not observed in ORF7-KO-iSLK cells (Figs. 1B and C). From these data, it was clarified that ORF7 plays an important role in the process of capsid formation.

**FIG 1.**
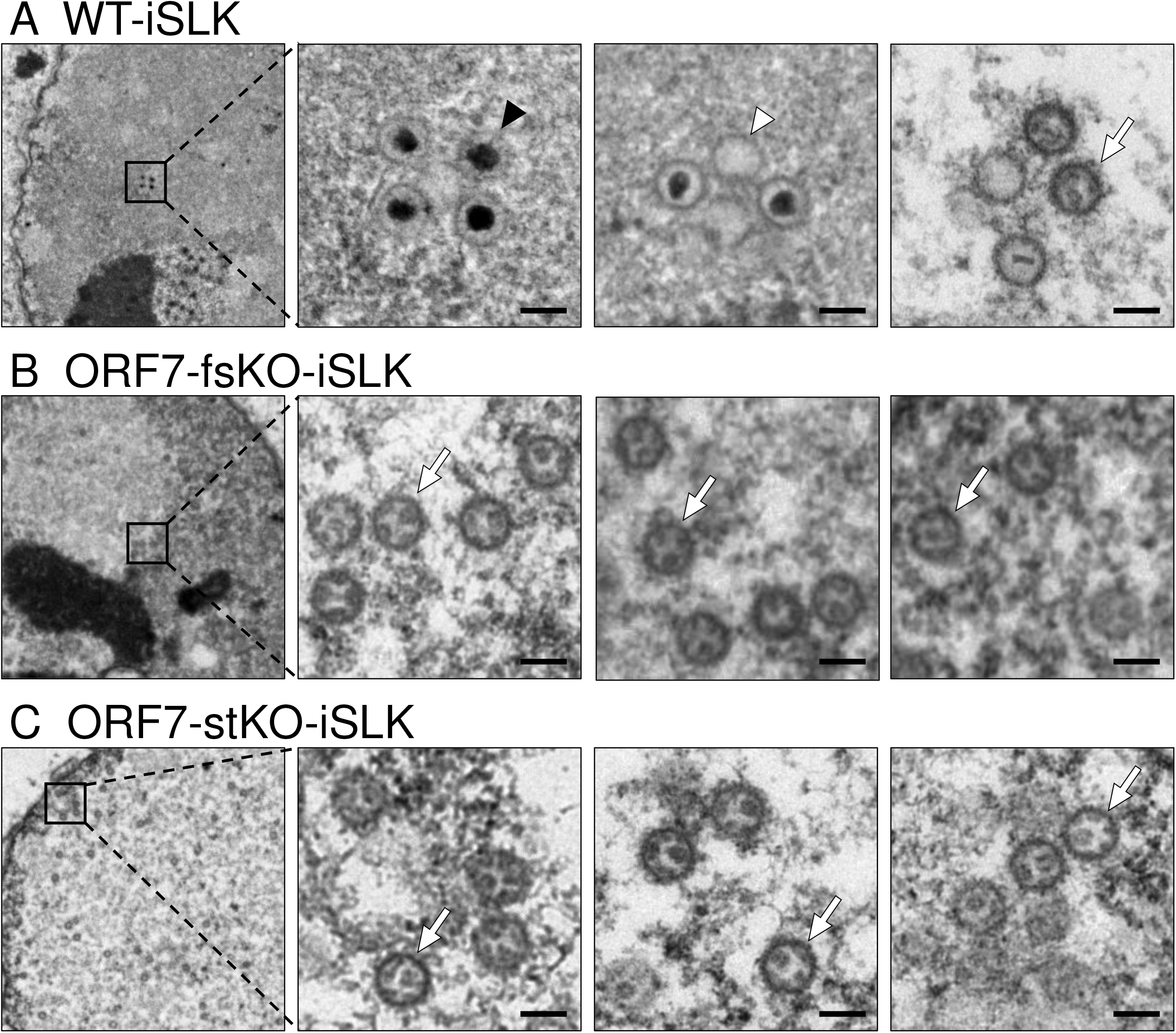
TEM images showing capsids produced from lytic-induced WT-KSHV- and ORF7-KO-KSHV-harboring iSLK cells. iSLK cells harboring WT or ORF7-KO KSHV were cultured for 72 h in medium containing Dox and NaB to induce the lytic phase. Capsid formation in nuclear (A) WT-iSLK, (B) frameshift-induced ORF7 KO (ORF7-fsKO-iSLK), and (C) stop codon-induced ORF7 KO (ORF7-stKO-iSLK) cells as observed by TEM. The black and white arrowheads indicate capsids and A-capsids, respectively, and the white arrows indicate soccer ball-like capsids. The scale bars represent a length of 100 nm.

### ORF7 is required for the cleavage of the KSHV TR during the lytic state

Since incomplete capsids (i.e., soccer ball-like capsids) were formed in the nuclei of iSLK cells harboring ORF7-KO KSHV during lytic infection, it was clear that ORF7 plays an important role in KSHV capsid formation. Next, we asked whether the KSHV ORF7 cleaves the KSHV genome, thus contributing to KSHV terminase activity. To address this question, the cleavage of replicated KSHV DNA containing tandemly repeated genomes was detected by Southern blotting according to the method established by Glaunsinger *et al.* (27, 28) with slight modifications (Fig. 2A). In this assay, a DIG-labeled one unit TR was used as the DNA probe to detect KSHV TRs. Lytic-induced cells treated with Dox and NaB were cultured for 72 h, and cellular and viral genomic DNA was purified and digested with restriction enzymes (EcoRI and SalI). Subsequently, agarose gel electrophoresis was performed (Fig. 2B). In the nucleus of an infected cell, one KSHV genome containing approximately 96 ORFs is circularized via binding of the TRs located at the 5’- and 3’-ends of the cording region. The TR region is comprised of approximately 40 TRs, which contain several viral terminase cleavage sites. In the lytic stage, KSHV genomes are replicated as a linear form containing tandemly repeating genomes, and each genomic unit is sequentially connected through a TR region. The KSHV terminase complex cleaves the DNA at the TR sites within the replicated KSHV genome and contributes to the production of an appropriate genome unit. EcoRI and SalI digest the cellular DNA and ORFs of the KSHV genome (BAC16), but not the KSHV TRs. The separated DNA was transferred to a nylon membrane and the TRs were detected by Southern blotting. Uncleaved TRs from BAC16 were detected in all samples: WT-BAC16- (WT-iSLK), ORF7-fsKO-BAC16- (ORF7-fsKO-iSLK), ORF7-stKO-BAC16-(ORF7-stKO-iSLK), and ORF7 revertant-BAC16-(ORF7-fsKO Rev-iSLK and ORF7-stKO Rev-iSLK) harboring iSLK cells. Cleaved TRs were detected in WT-iSLK, ORF7-fsKO Rev-iSLK, and ORF7-stKO Rev-iSLK. However, cleaved TRs were not detected in ORF7-fsKO-iSLK and ORF7-stKO-iSLK (Fig. 2B). These banding patterns of cleaved TR regions were in agreement with previous data which characterized an ORF68-KO reported by Glaunsinger *et al.* (27, 28). Our data indicated that ORF7 plays a critical role in TR cleavage during lytic infection-induced viral DNA replication.

**FIG 2.**
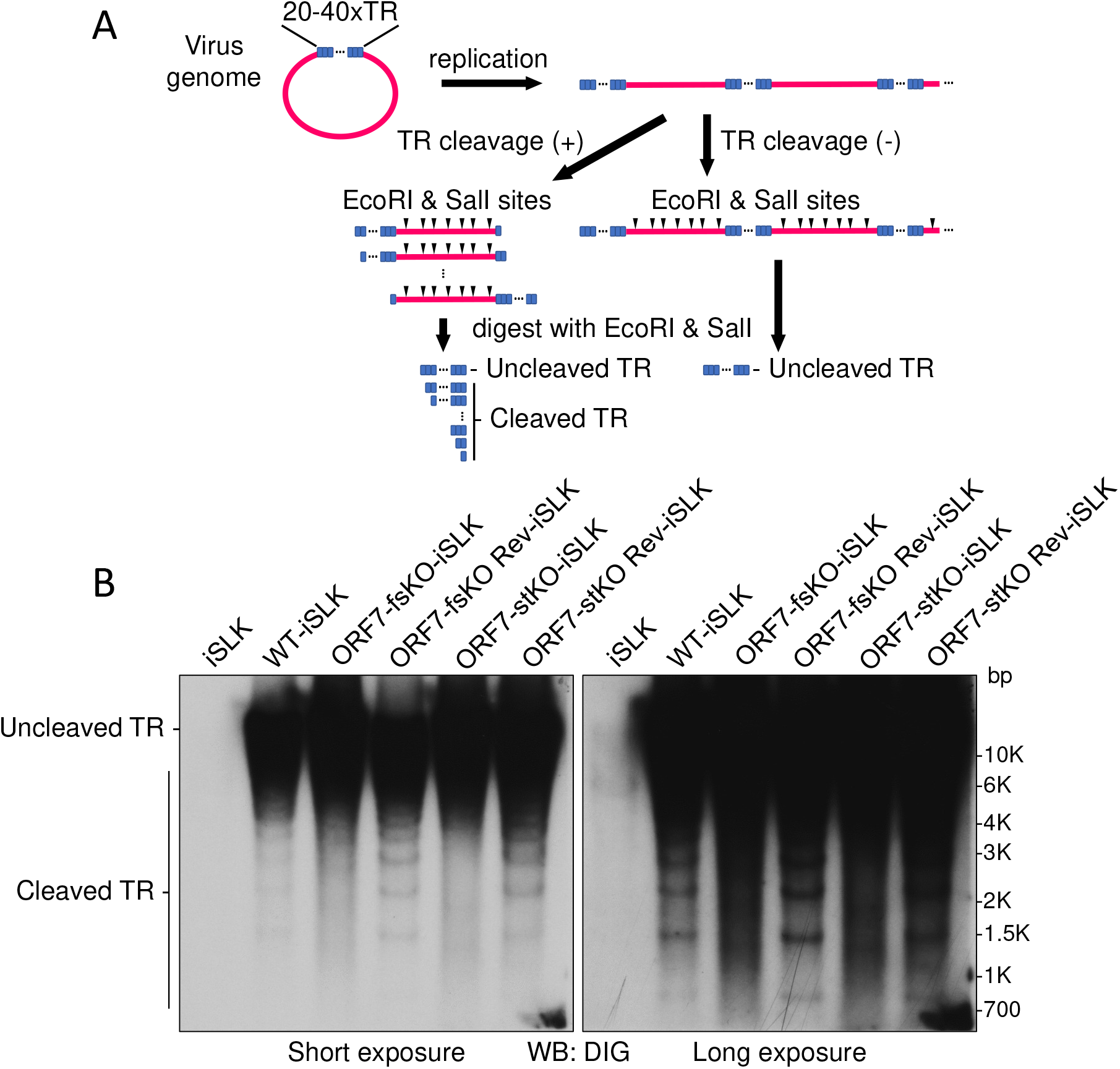
ORF7-KO-KSHV exhibits a genome cleavage defect. (A) Schematic diagram showing the Southern blotting procedure for detection of viral genome cleavage at the TR. Replicated viral genome, experimental procedure, and the DNA cleavage pattern with/or without TR cleavage. (B) Southern blotting of the EcoRI- and SalI-digested genomic DNA purified from Dox- and NaB-treated iSLK cells harboring WT-KSHV (WT-iSLK), ORF7-KO-KSHVs (ORF7-fsKO-iSLK and ORF7-stKO-iSLK), and the revertant-KSHVs (ORF7-fsKO Rev-iSLK and ORF7-stKO Rev-iSLK). The digested DNA was detected with a DIG-labeled TR probe. Left panel: short exposure, right panel: long exposure.

### ORF7 and ORF17 DKO KSHV produces procapsids possessing a circular inner scaffold, similarly to ORF17 KO KSHV

We previously showed that the KSHV protease ORF17 was involved in capsid maturation as a viral protease which digests scaffold proteins, resulting in scaffold shell disruption. Moreover, either complete KO of ORF17 or a mutant deficient in ORF17 protease activity induced capsid formation arrest between the procapsid and B-capsid stages (20). Here, we found that ORF7 possessed KSHV genome TR cleavage activity (Fig. 2B). The herpesvirus terminase complex contributes to viral genome cleavage and packaging of the cleaved genomes into immature capsids. This immature capsid is thought to be assembled after the formation of procapsids or internal scaffold shell-disrupted procapsids (29, 30). However, there are no reports describing which event (i.e., ORF7-mediated genome cleavage or ORF17-mediated internal scaffold disruption) occurs earlier during capsid formation. Thus, in order to determine the order of ORF7- and ORF17-mediated effects on capsid assembly, we constructed ORF7 and ORF17 DKO (designated as ORF7&17-DKO) KSHVs. We then compared viral events including capsid formation among WT, ORF7-KO, ORF17-KO, and ORF7&17-DKO KSHVs. If ORF7 acts earlier than ORF17 in capsid formation, then the ORF7&17-DKO KSHV should have the same phenotype as ORF7-KO KSHV. On the other hand, if ORF17 acts earlier than ORF7, then the ORF7&17-DKO KSHV should present the same phenotype as ORF17-KO KSHV. Thus, we constructed ORF7&17-DKO-BAC16 from ORF7-fsKO-BAC16 by insertion of the ORF17 KO frame shift mutation (which has been previously described) (20) and confirmed the DKO mutation by Sanger sequencing (Fig. 3A). The ORF7&17-DKO-BAC16 clone was transfected into the iSLK cell line, and stable cells harboring the BAC16 were selected by hygromycin B. Next, the Dox-inducible iSLK harboring ORF7&17-DKO-BAC16 (ORF7&17-DKO-iSLK) polyclonal cell line was established. We evaluated capsid formation in lytic-induced ORF7&17-DKO-iSLK cells by observation of capsid morphology using TEM. The WT-iSLK and ORF7&17-DKO-iSLK were treated with Dox and NaB for lytic induction, and cells were fixed to observe capsid formation. In the nucleus of WT-iSLK cells (Fig. 3B), C-capsids were predominant, and a few A- and B-capsids were also detected as shown in Fig. 1A. In contrast, B-like capsids having an internal spherical scaffold were mainly detected in ORF7&17-DKO-iSLK cells; however, soccer ball-like capsids were not observed (Fig. 3C). These B-like capsids observed in ORF7&17-DKO-iSLK cells are consistent with procapsids observed in ORF17 KO KSHV-harboring iSLK cells (20). On the other hand, ORF7 single-KO KSHV produced soccer ball-like capsids, a type of B-capsid possessing telstar pattern internal structures (Figs. 1B and C). These results indicated that in the KSHV capsid formation process, the soccer ball-like capsids observed in ORF7-KO KSHVs are assembled after the formation of procapsids and B-capsids. Moreover, ORF7 acts on capsid formation after ORF17 (Fig. 3D).

**FIG 3.**
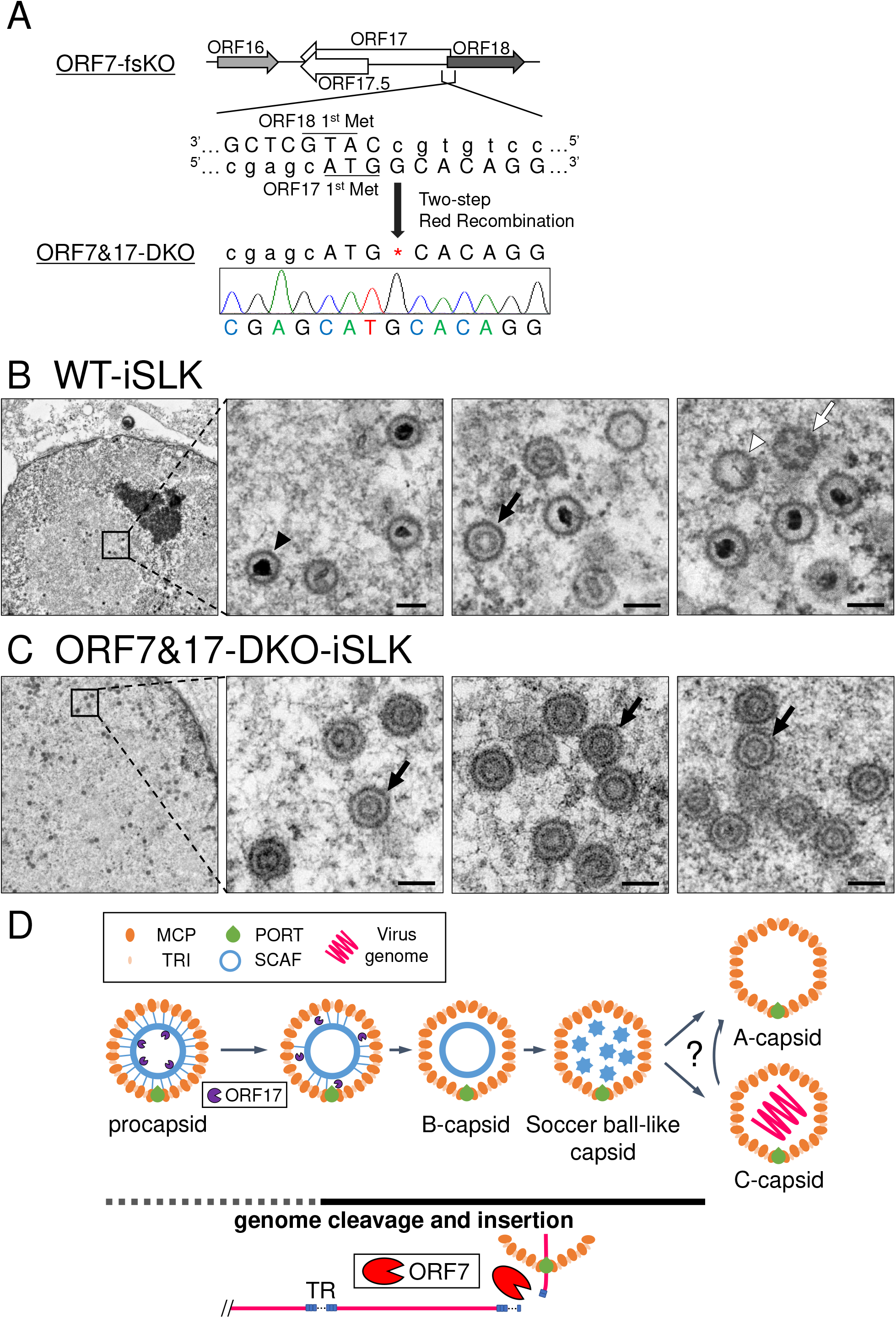
Construction of ORF7&17-DKO-KSHV, and TEM images of capsids produced from ORF7&17-DKO-KSHV. (A) Schematic illustration of the genetic locus of ORF17 and the mutation added to ORF7-fsKO-BAC16 for construction of ORF7&17-DKO-BAC16. The neighboring DNA sequence of the additional mutation in ORF7&17-DKO-BAC16 was confirmed by Sanger sequencing. (B,C) TEM images of capsids produced from WT-KSHV and ORF7&17-DKO-KSHV. iSLK cells harboring WT-KSHV (WT-iSLK) or ORF7&17-DKO-KSHV (ORF7&17-DKO-iSLK) were cultured for 72 h in medium containing Dox and NaB for induction of the lytic phase. Capsid formation in nuclei from (B) WT-iSLK cells and (C) ORF7&17-DKO-iSLK cells were observed by TEM. The black arrowheads, white arrowheads, black arrows, and white arrows indicate C-capsids, A-capsids, B-capsids, and soccer ball-like capsids, respectively. The scale bars represent a length of 100 nm. (D) Overview of the predictive KSHV capsid formation process including ORF7, ORF17 and soccer ball-like capsids. The predictive period of KSHV genome cleavage and genome insertion into capsids is represented by a bold line. MCP: major capsid protein, TRI: triplex, PORT: portal protein, SCAF: scaffold protein.

### Virological analysis of ORF7&17-DKO-KSHV

ORF7-KO induced the formation of KSHV soccer ball-like capsids (Figs. 1B and C), whereas ORF17-KO and ORF7&ORF17-DKO induced procapsids harboring an internal spherical scaffold, which resembled B-capsids (Fig. 3C) (20). In order to obtain further information regarding the viral properties of ORF7-KO-KSHV and ORF7&ORF17-DKO-KSHV, we evaluated viral genome replication, virion production, and viral transcription in the mutant KSHVs. The ORF7-fsKO-iSLK and ORF7&17-DKO-iSLK cells were treated with Dox and NaB to induce the lytic state, and the intracellular viral genome copy number was analyzed. There was no change in the amount of intracellular viral genomes among the tested cells (Fig. 4A). Next, we determined virus production and virus infectivity. The supernatants from lytic-induced cells were harvested, and encapsidated viral genome copies were measured by real-time PCR (qPCR). In addition, in order to measure the viral infectivity supernatants from lytic-induced cells were incubated with fresh HEK293T cells, and GFP-positive cells were analyzed by flow cytometry. The virus production derived from ORF7&17-DKO-iSLK cells significantly decreased compared to the amount of virus derived from WT-iSLK cells. This decrease was similar to the viral levels obtained with ORF7-fsKO-iSLK cells (Fig. 4B). The infectivity of ORF7&17-DKO-iSLK and ORF7-fsKO-iSLK was drastically reduced to nearly undetectable levels compared to the infectivity derived from WT-iSLK cells (Fig. 4C). As for viral transcription, mRNA expression levels of immediate early (IE) (ORF16), early (E) (ORF46 and ORF47), and late (L) (K8.1) genes in lytic-induced WT-iSLK, ORF7-fsKO-iSLK, and ORF7&17-DKO-iSLK were evaluated by real-time reverse-transcription PCR (qRT-PCR). Similar to viral genome replication, the expression levels of IE, E, and L genes were comparable among all of the tested cells (Fig. 4D), which indicated that KO of ORF7, and KO of both ORF7 and ORF17 do not interfere with viral gene transcription. We previously reported that ORF17-KO-iSLK cells had similar viral genome replication activity compared to WT-iSLK cells but was deficient in virion production (20). Thus, ORF7&17-DKO KSHV showed the same properties as ORF7-KO KSHV in viral genome replication and infectious virus production. These phenotypes are fully consistent with those of ORF17-KO KSHV. Our data revealed that ORF7-KO KSHV produced soccer ball-like capsids (Figs. 1B and C) and failed to produce infectious virus (Figs. 4B and C). Moreover, ORF7 acted on capsid formation after ORF17-mediated effects on capsid assembly (Fig. 3). Therefore, ORF7-deficent KSHV may induce capsid formation arrest between the B-capsid and mature C-capsid stages during the lytic phase. Meanwhile, it is known that viral transcription and viral genome replication occur independently of capsid formation. Therefore, it was not surprising that the KO of ORF7, ORF17, and both ORF7&17 did not impact viral transcription and viral genome replication.

**FIG 4.**
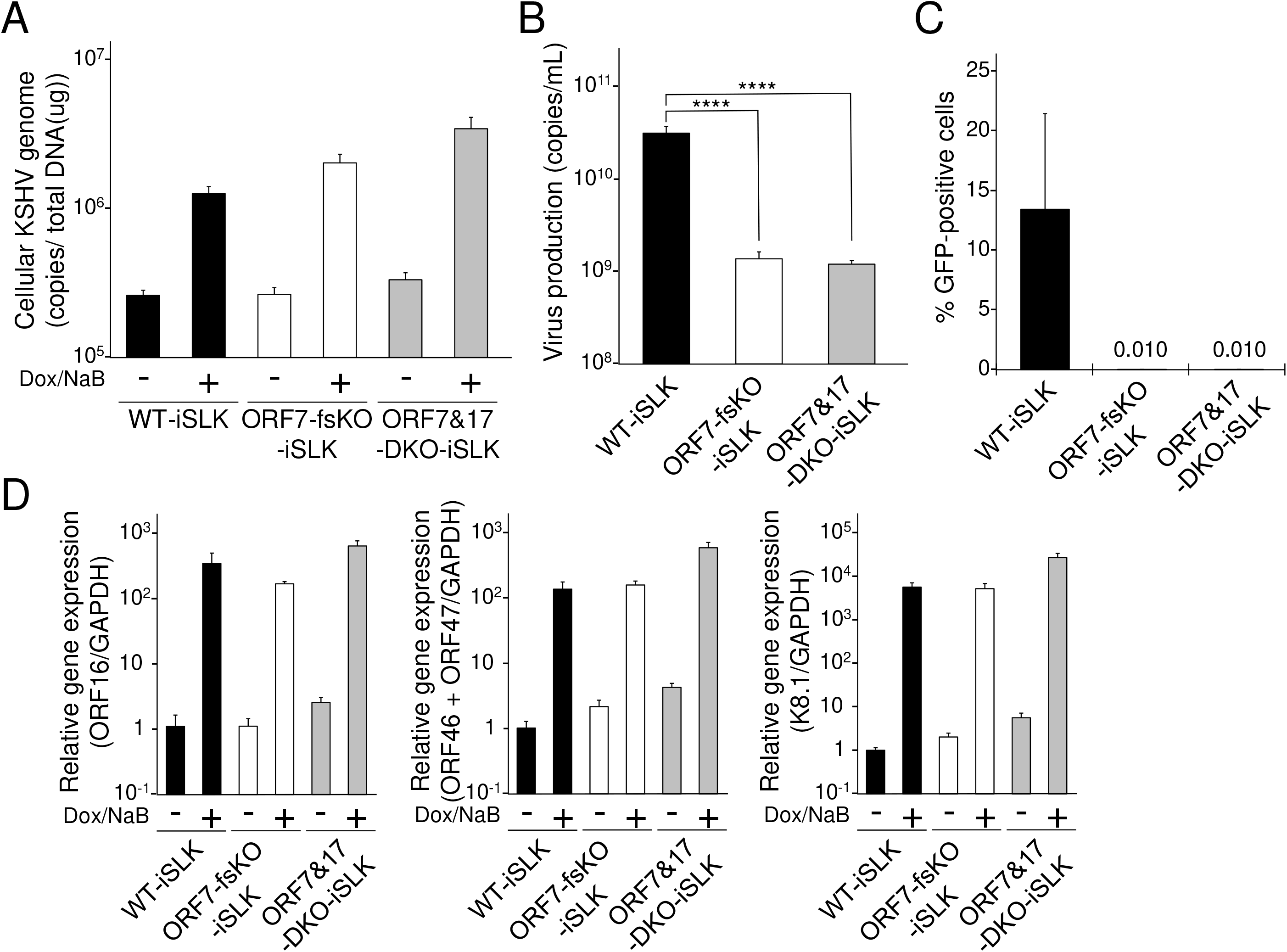
Characterization of ORF7-KO-KSHV and ORF7&17-DKO-KSHV. (A) Quantification of intracellular viral DNA (genome replication). WT-iSLK, ORF7-fsKO-iSLK, and ORF7&17-DKO-iSLK cells were treated with Dox and NaB to induce the lytic phase. Next, the intracellular viral DNA levels were measured using qPCR and were normalized to the total amount of DNA. (B) Quantification of extracellular encapsidated KSHV genomes. WT-iSLK, ORF7-fsKO-iSLK, and ORF7&17-DKO-iSLK cells were treated with Dox and NaB to induce the lytic phase and the culture supernatants were harvested. The encapsidated KSHV genome levels in the culture supernatants were measured using qPCR. ****P <0.001. (C) Measurement of infectious virus. WT-iSLK, ORF7-fsKO-iSLK, and ORF7&17-DKO-iSLK cells were treated with Dox and NaB to induce the lytic phase and the culture supernatants were harvested. Progeny virions from the cell supernatants were used to infect fresh HEK293T cells, and GFP-positive cells were counted by flow cytometry at 24h post-infection. (D) Analysis of mRNA expression of KSHV lytic genes: IE, ORF16; E, ORF46 and ORF47; and L, K8.1. Total RNA was purified from lytic-induced iSLK cells andwas subjected to qRT-PCR. The evaluated lytic gene mRNA levels were normalized to the GAPDH mRNA levels. The values obtained from Dox- and NaB-untreated WT-iSLK cells were defined as 1.0.

In order to evaluate the importance of both ORF7 and ORF17 in virus production, we analyzed whether exogenous ORF7 and ORF17 could rescue the infectious virus production in ORF7&17-DKO-iSLK cells. ORF7&17-DKO-iSLK cells were transfected with an empty plasmid, ORF7-2xS expressing plasmid alone (ORF7-2S), 3xFLAG-ORF17 expressing plasmid alone (3F-ORF17), or both ORF7-2xS (ORF7-2S) and 3xFLAG-ORF17 (3F-ORF17) expressing plasmids. The transfected cells were treated with Dox and NaB to induce the lytic phase. Subsequently, culture supernatants were subjected to qPCR to measure virus production. Virus production in ORF7&17-DKO-iSLK cells recovered significantly when ORF7-2xS (ORF7-2S) and 3xFLAG-ORF17 (3F-ORF17) were exogenously co-expressed (Fig. 5A). The protein expression of ORF7-2xS (ORF7-2S) and 3xFLAG-ORF17 (3F-ORF17) were confirmed by Western blotting (the lower panels of Fig. 5A). In addition to measuring viral DNA, we also evaluated the virus titers in the culture supernatants. In order to evaluate infectious virion production, the culture medium was added to uninfected HEK293T cells, and the numbers of GFP-positive cells were counted by flow cytometry. We found that the infectivity of the virions produced from ORF7&17-DKO-iSLK cells were recovered by exogenous ORF7-2xS (ORF7-2S) and 3xFLAG-ORF17 (3F-ORF17) co-expression (Fig. 5B). These complementation assays that evaluated the effects of exogenous ORF expression on virus production and infectivity indicated that ORF7 and ORF17 play crucial roles in virus production.

**FIG 5.**
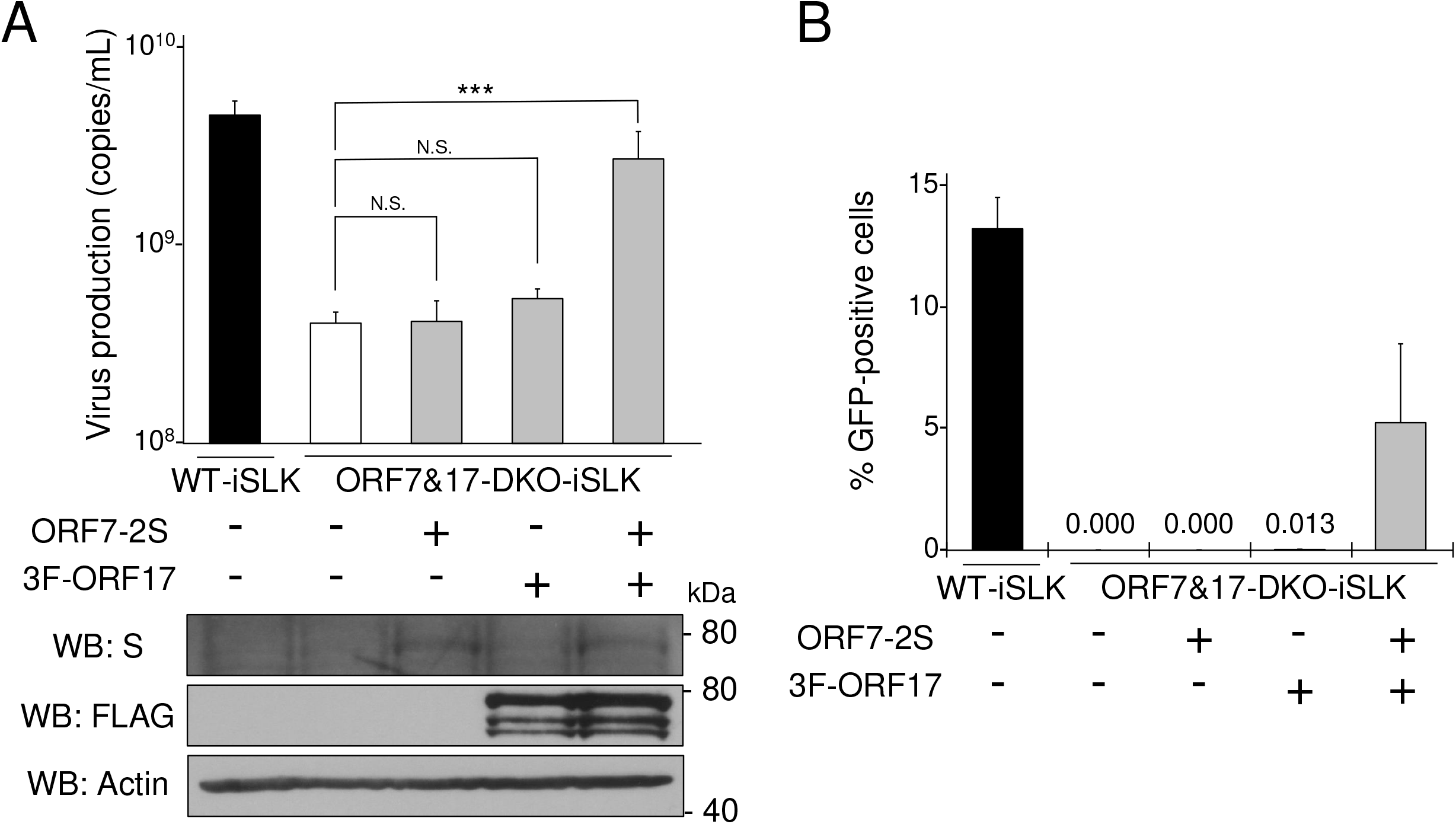
A decrease in virus production in ORF7&17-DKO-iSLK cells was overcome by exogenous expression of both ORF7 and ORF17. (A) Exogenous expression of both ORF7 and ORF17 rescued the suppression of extracellular KSHV levels observed in ORF7&17-DKO-iSLK cells. ORF7&17-DKO-iSLK cells were transiently transfected with ORF7-2xS (ORF7-2S) alone, 3xFLAG-ORF17 (3F-ORF17) alone, or both ORF7-2xS (ORF7-2S) and 3xFLAG-ORF17 (3F-ORF17) plasmids and were simultaneously cultured with medium containing Dox and NaB for 72 h to induce the lytic phase. Encapsidated KSHV genomes in the culture supernatant were quantitated using qPCR. Exogenous ORF7 and ORF17 expression were confirmed by Western blotting (WB) using anti-S (S) and anti-FLAG (FLAG) antibodies. Antibodies directed against β-actin (Actin) were used as a loading control. N.S., not significant (P>0.05). ***P<0.005. (B) Exogenous expression of both ORF7 and ORF17 rescued the suppression of infectious virus production observed in ORF7&17-DKO-iSLK cells. ORF7&17-DKO-iSLK cells were transiently transfected with ORF7-2xS (ORF7-2S) alone, 3xFLAG-ORF17 (3F-ORF17) alone, or both ORF7-2xS (ORF7-2S) and 3xFLAG-ORF17 (3F-ORF17) plasmids and were simultaneously cultured with medium containing Dox and NaB for 72 h to induce the lytic phase. The harvested culture supernatants were used to inoculate fresh HEK293T cells for infection. At 24 h post-infection, GFP-positive cells were counted by flow cytometry.

### The conserved zinc-finger motif of ORF7 is important for infectious KSHV production

The KSHV ORF7 harbors a putative zinc-finger motif (C182-X_2_-C185-X_22_-C208-X_1_-H210), which is highly conserved among other human herpesvirus homologous genes (25). The zinc-fingers, which binds zinc ion, are speculated to be formed by three C and one H (i.e., C182, C185, C208, and H210) within this motif (C182-H210). In order to investigate the importance of the ORF7 zinc-finger motif for infectious virus production, we generated a series of ORF7 zinc-finger mutants. These mutations consisted of amino acid substitutions of the three Cs and one H which form the zinc-finger motif. These mutations are shown in Fig. 6A. These mutants were subjected to a transient complementation assay using ORF7-fsKO-iSLK cells. Expression plasmids encoding 3xFLAG tagged ORF7 WT (ORF7 WT) or the ORF7 mutants [mC2 (C182A and C185A), mC1H1 (C208A and H210A), and mZf (C182A, C185A, C208A, and H210A)] (Fig. 6A) were transiently transfected into ORF7-fsKO-iSLK cells, and virus production was analyzed by qPCR and viral infectivity was assessed by flow cytometry. The ORF7 WT- or ORF7 mutant-transfected cells were cultured with medium containing Dox and NaB for lytic induction. The amount of virus production in the culture supernatant was determined by measurement of viral DNA using qPCR. Virus production was efficiently recovered by ORF7 WT expression; however, empty vector and all of the zinc-finger motif mutants (mC2, mC1H1, and mZf) did not induce virus production recovery (Fig. 6B). The protein expression of ORF7 WT and the ORF7 mutants were confirmed by Western blotting (the lower panels of Fig. 6B). Next, the amount of infectious virus in the culture supernatant, prepared using the same method as described for Fig. 6B, was determined. The culture medium was added to fresh HEK293T cells, and the number of GFP-positive cells were counted by flow cytometry to evaluate the infectious virion production. Our results agreed with the qPCR data (Fig. 6B). Specifically, the infectivity of the virions produced from ORF7-fsKO-iSLK cells were recovered by ORF7 WT, whereas all of the ORF7 zinc-finger motif mutants did not facilitate the recovery of viral infectivity (Fig. 6C). These results indicated that the zinc-finger motif of ORF7 is essential for the production of infectious KSHV.

**FIG 6.**
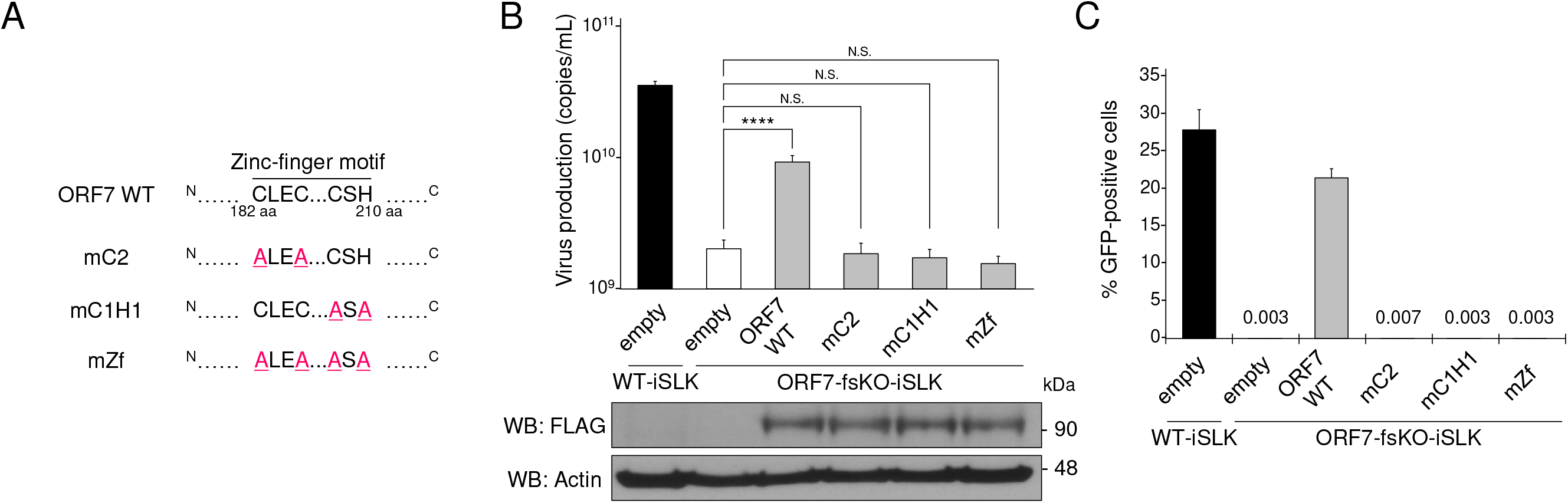
The conserved zinc-finger motif within ORF7 is essential for the recovery of suppressed virus production in ORF7-fsKO-iSLK cells. (A) The zinc-finger of ORF7, which binds to zinc ion, is speculated to be formed by three Cs and one H (i.e., C182, C185, C208, and H210) within the zinc-finger motif (C182-H210). Three expression plasmids encoding ORF7 zinc-finger mutants, along with a 3xFLAG tag were constructed. These mutations comprised the following amino acid substitutions: mC2 (C182A, C185A), mC1H1 (C208A, H210A), and mZf (C182A, C185A, C208A, H210A). (B) The effects of exogenous ORF7 zinc-finger mutants on suppressed extracellular encapsidated viral genome levels in ORF7-fsKO-iSLK cells. ORF7-fsKO-iSLK cells were transfected with ORF7 WT or ORF7 zinc-finger mutant plasmids and were simultaneously cultured with Dox and NaB for 72 h to induce the lytic phase. The culture supernatants were harvested and the levels of encapsidated KSHV genomes were determined by qPCR. In order to confirm the expression of ORF7 WT or the ORF7 mutants transfected and lytic-induced iSLK cells were lysed and subjected to Western blotting with anti-FLAG antibody (FLAG). Antibodies directed against β-actin (Actin) were used as a loading control. N.S., not significant (P>0.05). ****P<0.001. (C) The effects of exogenous ORF7 zinc-finger mutants on suppressed infectious virus production in ORF7-fsKO-iSLK cells. ORF7-fsKO-iSLK cells were transfected with ORF7 WT or the ORF7 mutants and were simultaneously cultured with Dox and NaB for 72 h to induce the lytic phase. The culture supernatants were harvested and were inoculated into fresh HEK293T cells to monitor infectivity. At 24 h post-infection, GFP-positive cells were counted by flow cytometry.

### Virological analysis of ORF7 zinc-finger motif mutant KSHV

In order to further explore whether the zinc-finger motif of ORF7 is important for virus production, we constructed ORF7 mC1H1 (C208A, H210A) BAC16. Since the mC1H1 mutation requires the narrowest mutation region compared to the other two mutations, the mC1H1 mutation was introduced into WT-BAC16 to generate an ORF7 zinc-finger motif mutant. To construct ORF7 mC1H1-BAC16, the alanine mutations (C208A, H210A) were introduced into WT-BAC16 using the same method (Fig. 3A) described for the construction of ORF7&17-DKO-BAC16. The introduced mutations were confirmed by Sanger sequencing (Fig. 7A). Complementation assays (Figs. 6B and C) revealed that the reduced levels of infectious virus in ORF7-fsKO-iSLK cells was not recovered by any of the three ORF7 mutants (mC2, mC1H1, and mZf). In contrast, ORF7 WT exhibited robust recovery. The ORF7 mC1H1-BAC16 clone was transfected into the iSLK cell line, and stable cells harboring the ORF7 mC1H1-BAC16 were selected with hygromycin B for the establishment of a Dox-inducible ORF7 mC1H1-iSLK polyclonal cell line. In order to characterize ORF7 mC1H1-BAC16, we measured the lytic gene mRNA expression levels, replication of cellular KSHV genome copies, and infectious virus production in ORF7 mC1H1-iSLK cells by the same methods described for Fig. 4. There were no marked changes in both lytic gene expression and intracellular viral genome copy number in ORF7 mC1H1-iSLK cells compared to WT-iSLK cells (Figs. 7B and C). On the other hand, in ORF7 mC1H1-iSLK cells, virus production was significantly suppressed, and the infectivity was abolished compared to WT-iSLK cells (Figs. 7D and E). Thus, our reverse genetic approach established that the zinc-finger motif of ORF7 is essential for virion production during KSHV lytic infection.

**FIG 7.**
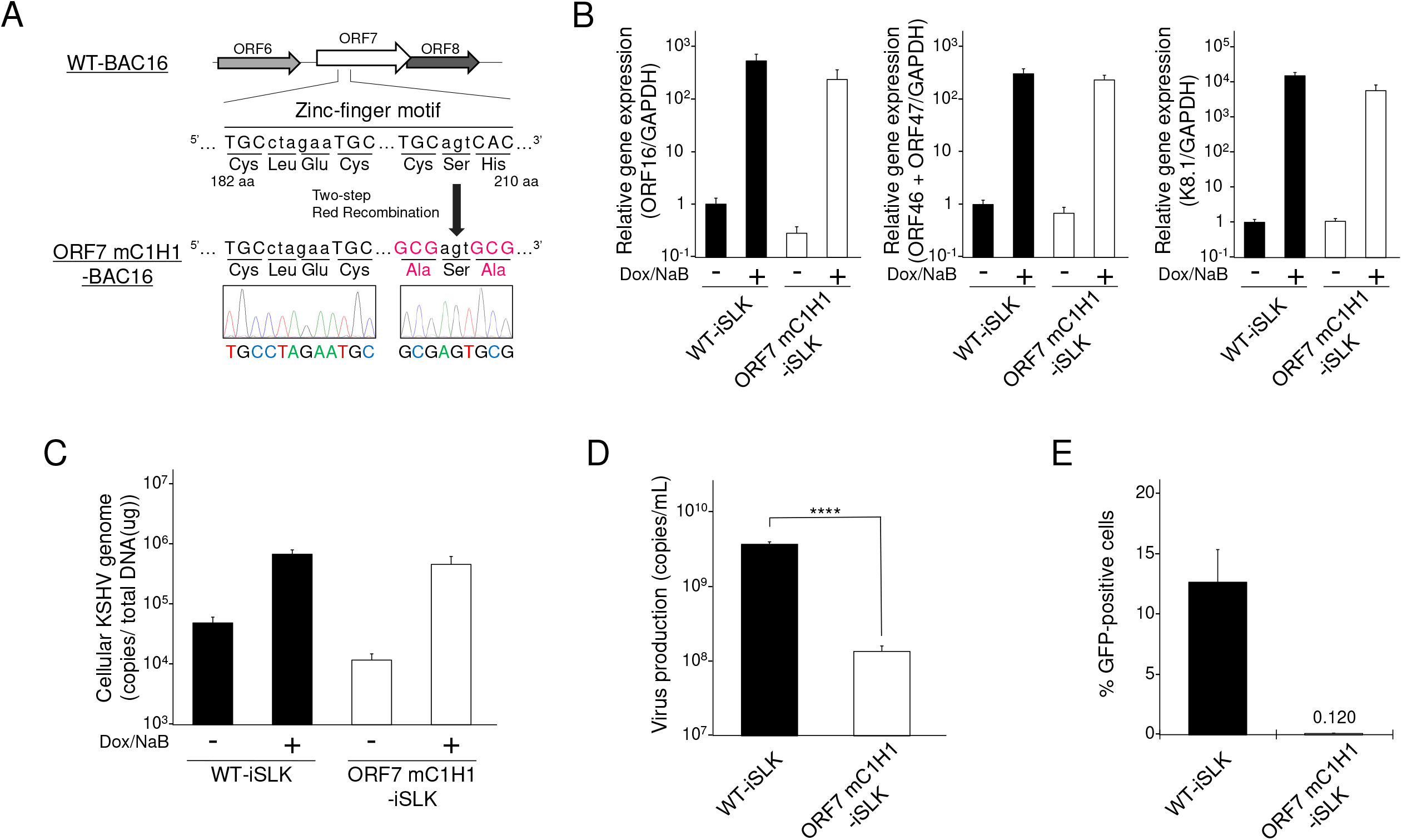
Construction and characterization of ORF7 mC1H1 KSHV. (A) Schematic illustration of the genetic locus of ORF7 and the mutation added to WT-BAC16 for construction of ORF7 mC1H1-BAC16. The neighboring DNA sequence of the additional mutation in ORF7 mC1H1-BAC16 was confirmed by Sanger sequencing. (B) Measurement of mRNA expression levels of viral lytic genes: IE, ORF16; E, ORF46 and ORF47; L, K8.1 in lytic-induced WT-iSLK and ORF7 mC1H1-iSLK cell lines. iSLK cells were treated with Dox and NaB for 72 h to induce the lytic phase, and total RNA was subjected to qRT-PCR. The mRNA levels of the tested lytic genes were normalized to the GAPDH mRNA levels. The values obtained from Dox- and NaB-untreated WT-iSLK cells were defined as 1.0. (C) Quantification of intracellular viral DNA (genome replication). iSLK cells were treated with Dox and NaB to induce the lytic phase. The intracellular viral DNA levels were measured using qPCR and were normalized to the total amount of DNA. (D) Quantification of extracellular encapsidated KSHV genome levels. iSLK cells were treated with Dox and NaB to induce the lytic phase. The culture supernatants were harvested and were assayed using qPCR to quantitate the encapsidated KSHV genome levels. ****P <0.001. (E) Measurement of infectious virus released from lytic-induced iSLK cells. iSLK cells were treated with Dox and NaB to induce the lytic phase and the culture supernatants were harvested. Progeny virions in the supernatants were used to infect fresh HEK293T cells, and GFP-positive cells were counted by flow cytometry at 24 h post-infection.

Finally, we asked whether the ORF7 zinc-finger motif was required for cleavage of the KSHV TR during lytic infection in ORF7 mC1H1-iSLK cells. The contribution of the ORF7 zinc-finger motif to TR cleavage activity in BAC16 was examined by the same Southern blotting method as described for Fig. 2. Uncleaved TRs were detected in all of the tested KSHV-(WT-, ORF7-fsKO-, ORF7-stKO-, and ORF7 mC1H1-BAC16) harboring cells. DNA fragments from cleaved TRs were detected in WT-iSLK cells but were not detected in ORF7-fsKO-iSLK and ORF7-stKO-iSLK cells (Fig. 8). Interestingly, two aberrantly-sized fragments, that differed from the banding pattern observed in WT-iSLK cells, were detected in the ORF7 mC1H1-iSLK cells (Fig. 8). These results suggested that the ORF7 zinc-finger motif regulates cleavage (or cleavage length) at the TR site. Thus, these data demonstrated that the zinc-finger motif of ORF7 is necessary for proper KSHV TR cleavage. In addition, ORF7 may function as a subunit of the KSHV terminase complex that cleaves the TRs on KSHV genomes during lytic infection.

**FIG 8.**
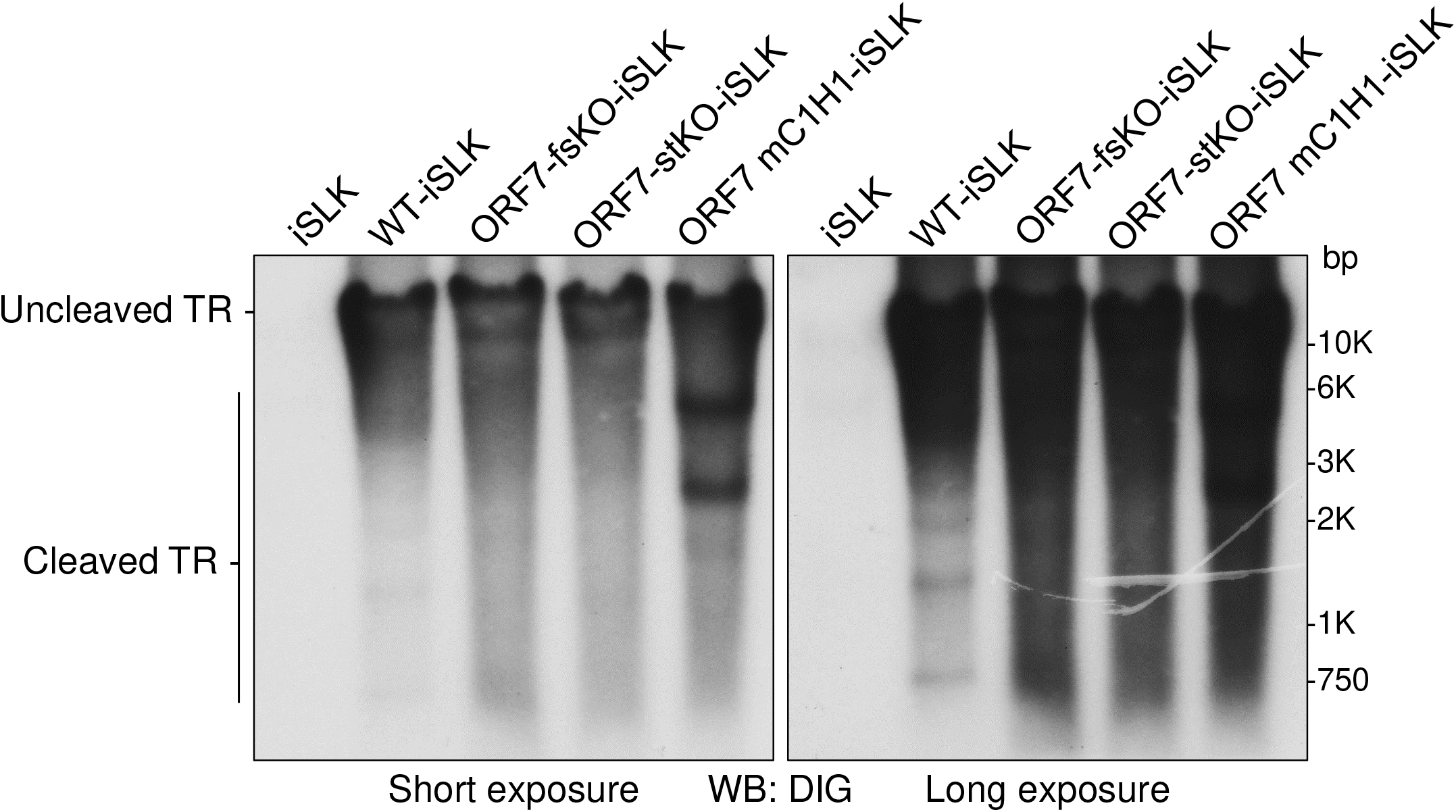
ORF7 mC1H1 KSHV failed to cleave the KSHV TR appropriately. Southern blotting of the EcoRI- and SalI-digested genomic DNA purified from Dox- and NaB-treated iSLK cells harboring WT-KSHV (WT-iSLK), ORF7-KO-KSHV (ORF-fsKO-iSLK and ORF7-stKO-iSLK) as well as ORF7 mC1H1-KSHV (ORF7 mC1H1-iSLK). Digested DNA was detected with a DIG-labeled TR probe. Left panel: short exposure, right panel: long exposure.

## Discussion

In this study, we showed that KSHV ORF7 is required for viral DNA cleavage and packaging, which suggested that it is important for KSHV terminase function. In addition, TEM analysis using both ORF7 and ORF17-deficient BAC16 confirmed that ORF7 acts after ORF17 in the KSHV capsid formation process. It was also revealed that the putative zinc-finger motif of ORF7 is essential for infectious virus production and contributes to proper TR cleavage by KSHV terminase. To our knowledge, this is the first report that used ORF-KO KSHV to show that ORF7 is involved in capsid formation and cleavage of viral genomes.

Soccer ball-like capsids observed in ORF7-deficient BAC16-harboring cells (Figs. 1B and C) were also detected in WT-BAC16-harboring cells (Fig. 1A). Although there are several reports including electron microscopy images showing completed C-capsids mixed with a few soccer ball-like capsids of varicella-zoster virus (VZV) and KSHV (30, 31, 32, 33), there have been no reports showing that an ORF deficiency resulted in the production of soccer ball-like capsids in herpesviruses. The soccer ball-like capsids can be detected during normal capsid formation of WT-KSHV, although our data revealed that only ORF7-KO induced the production of soccer ball-like capsids. Thus, we hypothesize two distinct origins of the soccer ball-like capsids: i) the soccer ball-like capsid is an intermediate capsid formed during the normal capsid assembly process, and ORF7-KO induces the arrest of capsid formation, resulting in the production of soccer ball-like capsids; or ii) the soccer ball-like capsids are not an intermediate form but a by-product of unsuccessful capsid formation, and a small amount of soccer ball capsids are produced as a by-product even during normal capsid formation. Further research on the mechanism of soccer ball-like capsid production is required to distinguish between these two hypotheses.

HSV-1 UL28 and human cytomegalovirus (HCMV) UL56, which are homologs of KSHV ORF7, were reported to be required for both cleavage of viral DNA and virus production (24, 34). These findings are consistent with the results obtained in our study. Capsid formation was arrested at the B-capsid stage by UL28 disruption in HSV-1 and UL56 inhibition in HCMV (24, 34). In both cases, the internal scaffold shell in these B-capsids retained its spherical form (24, 34). In contrast to HSV-1 and HCMV, KSHV ORF7 disruption produced soccer ball-like capsids lacking a spherical scaffold shell structure and instead had several small circular deposits. Here, we would like to consider the assembly process and internal components of soccer ball-like capsids. ORF7&17 double deficiency (Fig. 3C) and ORF17 single deficiency (20) produced incomplete capsids harboring an internal spherical scaffold shell. In contrast, ORF7 single deficiency produced soccer ball-like capsids (Figs. 1B and C). These results indicated that soccer ball-like capsids are products derived from incomplete capsids (B-like capsids) observed in ORF7&17-DKO and ORF17-KO KSHV-harboring iSLK cells, because ORF7 acts on capsid formation after ORF17-mediated capsid processing. Thus, we hypothesize that the soccer ball-like capsid is a sub-type of B-capsid that forms after the collapse of the B-capsid spherical scaffold. Since ORF7 functions as a DNA cleavage enzyme in the terminase complex, ORF7-KO KSHV may not only fail to cleave the viral genome, but also fail to package the cleaved genome into the capsid. Thus, we propose that the soccer ball-like capsid retains the collapsed scaffold shell debris and does not include viral DNA. In KSHV, the B-capsid contains an ensemble of “B-capsid-like” particles with highly diverse internal structures rather than a structurally homogeneous group with spherical scaffolds (30). The variable scaffold structures of B-capsids comprise not only a spherical shell form but also an aggregate form, and a spherical shell and aggregates emerge as the same density in cryo-electron tomography analysis (30). Thus, we propose that in ORF7-KO KSHV, the packaging of DNA into a capsid fails, and the scaffold proteins accumulate in a capsid without elimination from the capsid. Subsequently, scaffold proteins form aggregates that appear as a soccer ball pattern.

It is unclear whether soccer ball-like capsids observed in ORF7-deficent KSHV are formed as an intermediate capsid in the normal C-capsid assembly process or if they are produced as a by-product by unsuccessful capsid formation. However, our data indicated that soccer ball-like capsids are produced from B-capsid-like particles, which contain an internal scaffold induced by disruption of a spherical scaffold. There are several reports describing KSHV B-capsid formation. It has been reported that KSHV B-capsids retaining a spherical scaffold were produced by the KO of either ORF43 or ORF68 (27, 35). ORF43 forms a portal in the viral capsid for insertion of viral DNA into capsids, and ORF68 binds DNA and contributes to DNA cleavage and packaging (27, 35). In fact, in ORF43 or ORF68 KO, the KSHV capsid formation process is arrested at the B-capsid stage. If soccer ball-like capsids are produced from B-capsids, the effects associated with ORF43 and/or ORF68 could be triggers for the transition from B-capsids to soccer ball-like capsids. Thus, we hypothesize that soccer ball-like capsid production is not derived from spontaneous collapse of the internal scaffold shell in B-capsids; rather, they are derived from the functions associated with ORF43 (DNA cleavage) or ORF68 (DNA packaging). It has been reported that six KSHV capsid proteins (ORF25, ORF62, ORF26, ORF17, ORF17.5, and ORF65) expressed in Sf9 cells self-assembled into angularized capsids containing a spherical internal structure. Thus, molecules other than these six ORFs may trigger the collapse or elimination of the internal scaffold shell in B-capsids (36). In addition, it has been reported that a KSHV ORF19-DQ (D173A and Q177A) mutant produced incomplete B-capsids (37). ORF19 is the cap protein of the portal within the capsid, and the ORF19-DQ mutant fails to self-assemble into a pentamer. Authors mentioned these incomplete capsids as B-capsids observed in ORF19-DQ mutant KSHV (37). However, these capsids resemble soccer ball-like capsids observed in the ORF7-KO KSHV. So far, there have been few information about soccer ball-like capsids. Therefore, it remains controversial whether the soccer ball-like capsid is a by-product of unsuccessful capsid formation or an intermediate capsid assembled during the normal capsid formation process. We need to clear this question in future studies.

KSHV ORF7 possesses a putative zinc-finger motif (C182-X_2_-C185-X_22_-C208-X_1_-H210), which is highly conserved among other human herpesviral homologous genes including HSV-1 UL28 and EBV BALF3. The conserved zinc-finger motifs of UL28 and BALF3 are important for viral DNA packaging and nuclease activity toward DNA genomes, respectively (21, 25). Our complementation assays with the conserved zinc-finger motif mutants of KSHV ORF7 showed that this motif is essential for infectious virus production (Figs. 6B and C). In addition, ORF7 zinc-finger motif mutant-KSHV (ORF7 mC1H1-BAC16) demonstrated that this motif is necessary for proper TR cleavage (Fig. 8). Uncleaved TRs were detected in WT-BAC16, ORF7-fsKO-BAC16, ORF7-stKO-BAC16, and ORF7 mC1H1-BAC16-harboring cells. WT-KSHV produced cleaved fragments from the BAC16 TR, whereas ORF7-KO-KSHV failed to produce cleaved fragments. Interestingly, the ORF7 mC1H1-KSHV produced two major aberrantly-sized fragments, which was dissimilar to the cleavage pattern of WT-KSHV. These data indicated that the ORF7 zinc-finger motif may distinguish the appropriate TR cleavage sites for the terminase, rather than regulating the nuclease activity of the terminase.

We found that KSHV ORF7 contributed to capsid formation as well as viral DNA cleavage, which is a main function of the viral terminase complex. This study provides the first evidence that ORF7 is necessary for KSHV terminase function. Since the machinery for viral genome cleavage and packaging into capsids by the viral terminase does not exist in the host cells, this machinery may be a promising target for the development of antiviral drugs with high virus specificity. We have previously reported that KSHV ORF7, ORF29, and ORF67.5 form a tripartite complex, in which ORF7 functions as a center hub molecule (26). Similarly, HSV-1 UL15 (KSHV ORF29 homolog), UL28 (KSHV ORF7 homolog), and UL33 (KSHV ORF67.5 homolog) form a tripartite complex, and hexameric ring of this tripartite complex becomes an HSV-1 terminase (22). Either UL15 or UL33 KO failed to cleave viral DNA and produced incomplete capsids containing a compact internal scaffold (38, 39). It was reported that the C-terminal exon region of KSHV ORF29 has nuclease activity (40). However, the contribution of ORF29 to viral replication has not yet been clarified. The role of KSHV ORF67.5 (HSV-1 UL33 homolog) in both viral replication and viral DNA cleavage remains unknown. The contributions of ORF29 and ORF67.5 to viral DNA cleavage activity and capsid formation needs to be investigated in future studies. Moreover, elucidation of the biological significance of the tripartite terminase complex composed of ORF7, ORF29, and ORF67.5, will further expand our understanding of KSHV genomic DNA processing and DNA packaging into capsids.

### Plasmids

The C-terminal 2xS-tagged and 3xFLAG-tagged ORF7 expression plasmids and the N-terminal 3xFLAG-tagged ORF17 expression plasmid have been previously described (20, 26). ORF7 mutants (mC2, mC1H1, and mZf) were amplified by conventional and overlap extension PCR utilizing KOD -Plus-Neo (TOYOBO, Osaka, Japan). The following PCR primers were used: S_EcoRI_ORF7, CATGAATTCATGGCAAAGGAACTGGCGGC; As_XbaI_ORF7, GTCTCTAGAGACCTGGGAGTCATTGTGGTTGC; S_ORF7_C182A_C185A, CCGCCCTAGAAGCCCTTCAGGAAGTGTGTCTG; As_ORF7_C182A_C185A, GAAGGGCTTCTAGGGCGGGCAGATTTGAGACATATAG; S_ORF7_C208A_H210A, CCGCAAGTGCAATATGTACCCCCGCATGCG; and As_ORF7_C208A_H210A, TATTGCACTTGCGGCCGTGTCTGGGAGC. The plasmid pCI-neo ORF7 WT-3xFLAG was used as the PCR template. The amplified inserts were cloned into the pCI-neo mammalian expression vector (Promega, WI, USA) with the DNA Ligation Kit, Mighty Mix (Takara Bio, Shiga, Japan). The insert sequences were verified by Sanger sequencing.

### Construction of ORF7&17-DKO-BAC16 and ORF7 mC1H1-BAC16

ORF7&17-DKO-BAC16 was generated from ORF7-fsKO-BAC16 using a two step markerless Red recombination system (26, 41, 42). ORF7 mC1H1 (C208A, H210A) mutated BAC16 was generated from WT-BAC16. The mutagenesis of these BAC clones were performed according to a previously described protocol (20), using the following mutagenesis primers: S_dORF17_fs_EP, GCTCAGCGGCTCGGTCTCACACACGTATTTTCCGAGCATGCACAGGGCCTGTACGTCGGATAGGGATAACAGGGTAATCGATTT; As_dORF17_fs_EP, GACACAACATCTACAAACCCTCCGACGTACAGGCCCTGTGCATGCTCGGAAAATACGTGTGCCAGTGTTACAACCAATTAACC; S_ORF7_mC1H1_EP, CCAGGGCACCAGTCTGCAGGCCATGCTCCCAGACACGGCCGCGAGTGCGATATGTACCCCCGCATGCGGTAGGGATAACAGGGTAATCGATTT; and As_ORF7_mC1H1_EP, AGAGGCCCCGGACAGGCTCACCGCATGCGGGGGTACATATCGCACTCGCGGCCGTGTCTGGGAGCATGGGCCAGTGTTACAACCAATTAACC. The BAC clones containing the ORF7 and ORF17 mutated sites as well as the ORF7 mC1H1 alanine mutation (accession number: GQ994935) were confirmed by Sanger sequencing.

### Establishment of tetracycline/doxycycline-inducible recombinant KSHV-expressing iSLK cells

In order to obtain efficient recombinant KSHV-producing cells, tetracycline/doxycycline (Dox)-inducible (Tet-On) RTA/OR F50-expressing iSLK cells were used as virus-producing cells (43, 44, 45). iSLK cells were cultured in Dulbecco Modified Eagle’s medium supplemented with 10% fetal bovine serum, 1 ug/mL of puromycin (InvivoGen, CA, USA), and 0.25 mg/mL of G418 (Nacalai Tesque Inc., Kyoto, Japan). Thirty-six ug of WT-BAC16 or its mutants (ORF7-fsKO-BAC16, ORF7-stKO-BAC16, ORF7-fsKO Rev-BAC16, ORF7-stKO Rev-BAC16, ORF7&17-DKO-BAC16, and ORF7 mC1H1-BAC16) were transfected into iSLK cells (1 x 10^6^ cells) using the calcium phosphate method. WT-BAC16 was a kind gift from Jae U. Jung and Dr. Kevin Brulois. ORF7-fsKO-BAC16, ORF7-stKO-BAC16, ORF7-fsKO Rev-BAC16, and ORF7-stKO Rev-BAC16 were previously constructed (26, 42). The transfected cells were selected under 1 mg/mL of hygromycin B (Wako, Osaka, Japan) to establish Dox-inducible recombinant KSHV-producing cell lines (WT-iSLK, ORF7-fsKO-iSLK, ORF7-stKO-iSLK, ORF7-fsKO Rev-iSLK, and ORF7-stKO Rev-iSLK, ORF7&17-DKO-iSLK, and ORF7 mC1H1-iSLK).

### Measurement of viral gene expression, viral DNA replication, encapsidated viral DNA, **and infectious virus**

iSLK cells harboring WT or each KSHV mutant were treated with 8 μg/mL of Dox and 1.5 mM of sodium butyrate (NaB) for 72 h to induce the lytic phase (43, 44, 45). For quantification of viral gene expression, lytic-induced cells (5 x 10^5^ cells on a 6-well plate) were washed with PBS and harvested with 500 uL of RNAiso Plus (Takara Bio). Total RNA was extracted according to the manufacturer’s protocol. cDNA was synthesized from 160 ng of total RNA using the ReverTra Ace® qPCR RT Master Mix (TOYOBO). In order to quantitate the levels of viral lytic mRNAs, qRT-PCR was conducted using the synthesized cDNAs as template.

For measurement of intracellular viral DNA replication, iSLK cells (3.5 x 10^4^ cells on a 48-well plate) were treated with Dox and NaB for 72 h and harvested. The viral genomic DNA was purified from harvested cells with a QIAamp DNA Blood Mini Kit (QIAGEN, CA, USA). The viral genome copy number was quantified by qPCR and normalized to the amount of total DNA.

For quantification of extracellular encapsidated viral DNA, iSLK cells (1.5 x 10^5^ cells on a 12-well plate) were treated with Dox and NaB for 72 h. The culture supernatants (400 uL) were harvested and centrifuged at 15,000 rpm for 10 min to remove debris. The supernatants (220 uL) were treated with DNase I (New England Biolabs, MA, USA), and encapsidated viral DNA was extracted from the supernatants (200 uL) using the QIAamp DNA Blood Mini Kit. The purified KSHV genomic DNA was quantitated using qPCR.

For measurement of infectious virus, iSLK cells (2 x 106 cells on a 10 cm dish) were treated with Dox and NaB for 72 h, and the culture supernatants and cells were collected. The supernatants and cells (2.5 mL) were centrifuged at 15,000 rpm for 10 min at room temperature. Next, the collected supernatant (2 mL) was mixed with detached HEK293T cells (7.5 x 10^5^ cells). Polybrene (8 ug/mL) (Sigma-Aldrich, MO, USA) was added to the cell mixtures, which were then placed into 6-well plates. The plates were centrifuged at 1,200 x g for 2 h at room temperature and incubated at 37°C with 5%CO_2_. After 24 h, GFP-positive cells were analyzed with a flow cytometer [FACSCalibur (Beckton Dickinson, CA, USA)] using the CellQuest Pro software (Beckton Dickinson, CA, USA).

### qRT-PCR and qPCR

All qRT-PCRs were performed using the THUNDERBIRD® Next SYBR® qPCR Mix (TOYOBO) (43, 44). The following primers were used for quantification of viral gene expression: Fw-GAPDH, CATCAAGAAGGTGGTGAAGCAG; Rv-GAPDH, TGTCGCTGTTGAAGTCAGAGG; Fw-ORF16, AGATTTCACAGCACCACCGGTA; Rv-ORF16, CCCCAGTTCATGTTTCCATCGC; Fw-ORF47, CGATCCGAATCACTGCAACG; Rv-ORF47, CTGCTGCTTTTAGCCCGAG; Fw-K8.1, TCCCACGTATCGTTCGCATTTGG; and Rv-K8.1, GCGTCTCTTCCTCTAGTCGTTG. The relative mRNA expression levels were determined by ΔΔCt methods and were normalized to GAPDH mRNA levels. For measurement of KSHV genome copies, qPCR was conducted with the following KSHV-encoded ORF11-specific primers: KSHV ORF11-F, TTGACAACACGCACCGCAAG; and KSHV ORF11-R, AAAAATCAGCACGCTCGAGGAG.

### Complementation assay

The iSLK cells were transfected with each plasmid using ScreenFect A plus (Wako) according to the manufacturer’s instructions and the transfected cells were simultaneously cultured with medium containing Dox and NaB to induce the lytic state. After 72 h, the culture supernatants and cells were collected, and the amount of virus production and infectivity was assessed.

### Western blotting and antibodies

Western blotting was performed as described previously (46). Briefly, the cells were washed with PBS, lysed with SDS sample buffer, sonicated for 15 sec, boiled for 5 min, and subjected to SDS-PAGE and Western blotting. Anti-FLAG mouse mAb (M185) (MBL, Nagoya, Japan), horseradish peroxidase (HRP)-conjugated anti-S-tag rabbit pAb (ab18589) (Abcam, Cambridge, UK), and anti-β-actin mouse mAb (sc-69879) (Santa Cruz Biotechnology, TX, USA) were used as the primary antibodies. Anti-mouse IgG-HRP (NA931) (GE Healthcare, Buckinghamshire, UK) was used as the secondary antibody.

### Electron microscopy

iSLK cells (WT-iSLK, ORF7-fsKO-iSLK, ORF7-stKO-iSLK, and ORF7&17-DKO-iSLK) were cultured for 72 h in 8 μg/mL Dox and 1.5 mM NaB containing medium to induce the lytic phase. The cultured cells that were induced for virus production were washed with PBS and treated with a trypsin/EDTA solution for cell detachment. The samples were then fixed in 2% glutaraldehyde in PBS (pH 7.2) on ice for 2 h followed by incubation in 1% osmium tetroxide in PBS at 4°C for 1.5 h. The fixed samples were washed five times in PBS and dehydrated in ethanol using serially graded concentrations. The samples were then embedded in an epoxy resin (Nisshin EM Co., Ltd., Tokyo, Japan) and processed using a routine electron microscopy procedure. Ultrathin sections were prepared using a Porter Blum ultramicrotome (Reichert–Nissei ULTRACUT-N) (Leica, Wetzlar, Germany) and mounted on a copper grid (200 mesh) supported by a carbon-coated collodion film. The ultrathin sections were doubly stained with 4% (w/v) uranyl acetate and lead citrate for 10 min and 3 min, respectively. The sections were washed three times in distilled water after each staining procedure. All of the sections were observed under a Hitachi H-7800 transmission electron microscope.

### Southern blotting

For preparation of the DNA probe for Southern blotting, WT-BAC16 DNA was digested with NotI (TOYOBO) overnight and subjected to agarose gel electrophoresis to purify one unit of the TR sequence migrating near 800 bp. The extracted TR was cloned into a pCI-neo empty vector. The cloned sequence was confirmed to be a single copy of the KSHV TR by agarose gel electrophoresis and Sanger sequencing. Thus, the construct was referred to as the pCI-neo 1xKSHV TR (YI47). The DIG-labeled DNA probe for Southern blotting was prepared using the DIG-High Prime DNA Labeling and Detection Starter Kit II (Roche, Basel, Switzerland). As a template, we utilized the KSHV 1xTR (1 ug) obtained from NotI-digested pCI-neo 1xKSHV TR (YI47).

The lytic-induced iSLK cells (2 x 10^6^ cells on a 10 cm dish) were washed with PBS and suspended in TNE buffer [10 mM Tris-HCl (pH7.5), 100 mM NaCl, 1 mM EDTA]. In order to prepare the cell extract, SDS (0.3%) and proteinase K (100 ug/mL) were added to the cell suspension and the mixture was incubated overnight at 60°C. Subsequently, the cell extracts were incubated with RNase (100 ug/mL) at room temperature for 30 min. DNA was isolated from the cell extracts by phenol-chloroform extraction and ethanol precipitation. The pellets were then resuspended in 1xTE buffer. The DNA samples (10 ug each) were digested overnight with EcoRI (Takara Bio) and SalI (TOYOBO) followed by electrophoresis on a 0.7% agarose 1xTBE gel. As for Southern blotting (27, 28), the gel was denatured in alkaline denaturing buffer (0.5 N NaOH, 1.5 M NaCl) for 45 min and transferred to Hybond-N+ (Cytiva, Tokyo, Japan) by capillary action overnight in the presence of alkaline transfer buffer (0.4 N NaOH, 1 M NaCl). The transferred DNA was cross-linked to the membrane with a CL-1000 Ultraviolet Crosslinker (UVP, Cambridge, UK). The membrane was treated according to the DIG-High Prime DNA Labeling and Detection Starter Kit II (Roche, Basel, Switzerland) protocol. The membrane was hybridized at 42°C overnight in DIG Easy Hyb buffer with a DIG-labeled TR probe. Next, the membrane was washed two times with 2xSSC, 0.1% SDS at room temperature followed by an additional two washes with 0.5xSSC, 0.1% SDS at 68°C. The membrane was blocked, incubated with anti-DIG antibody, and detected by CSPD. The blot was then exposed to X-ray film (Fuji film, Tokyo, Japan).

### Statistics

The statistical significance was analyzed by one-way ANOVA followed by Dunnett’s test. The statistical significance was assessed using the GraphPad Prism 7 software (GraphPad Software, CA, USA).

## Acknowledgements

The KSHV BAC clone, BAC16, was a kind gift from Dr. Jae U. Jung (Cleveland Clinic Lerner Research Institute, USA). We thank Dr. Gregory A. Smith (Northwestern Univ., USA) for the *E. coli* strain GS1783, Yoshihiko Fujioka (Osaka Medical and Pharmaceutical Univ., Japan) for electron microscopy analysis, and Dr. Nikolaus Osterrieder (Cornell Univ., USA) for the plasmid pEP-KanS. Y.I. was supported by the Nagai Memorial Research Scholarship from the Pharmaceutical Society of Japan. M.F. was supported by a grant from the JSPS Grant-in-Aid for Scientific Research (18K0664).

## References

1. Chang Y, Cesarman E, Pessin MS, et al. Identification of herpesvirus-like DNA sequences in AIDS-associated Kaposi’s sarcoma. Science. 1994;266(5192):1865–1869. doi:10.1126/science.7997879

2. Nador RG, Cesarman E, Chadburn A, et al. Primary effusion lymphoma: a distinct clinicopathologic entity associated with the Kaposi’s sarcoma-associated herpes virus. Blood. 1996;88(2):645–656.

3. Soulier J, Grollet L, Oksenhendler E, et al. Kaposi’s sarcoma-associated herpesvirus-like DNA sequences in multicentric Castleman’s disease. Blood. 1995;86(4):1276–1280.

4. Russo JJ, Bohenzky RA, Chien MC, et al. Nucleotide sequence of the Kaposi sarcoma-associated herpesvirus (HHV8). Proc Natl Acad Sci U S A. 1996;93(25):14862–14867. doi:10.1073/pnas.93.25.14862

5. Uldrick TS, Wang V, O’Mahony D, et al. An interleukin-6-related systemic inflammatory syndrome in patients co-infected with Kaposi sarcoma-associated herpesvirus and HIV but without Multicentric Castleman disease. Clin Infect Dis. 2010;51(3):350–358. doi:10.1086/654798

6. Pellett P.E., Roizman B. Herpesviridae. In: Knipe D.M., Howley P.M., editors. Fields Virology. 6th ed. Lippincott Williams & Wilkins; Philadelphia, PA, USA: 2013. pp. 1802–1822.

7. Brown JC, Newcomb WW. Herpesvirus capsid assembly: insights from structural analysis. Curr Opin Virol. 2011;1(2):142–149. doi:10.1016/j.coviro.2011.06.003

8. B. Roizman, D.M. Knipe, R.J. Whitley Herpes simplex viruses D.M. Knipe, P.M. Howley (Eds.), Fields Virology (sixth ed.), Lippincott Williams & Wilkins (2013), pp. 1823–1897

9. Heming JD, Conway JF, Homa FL. Herpesvirus Capsid Assembly and DNA Packaging. Adv Anat Embryol Cell Biol. 2017;223:119–142. doi:10.1007/978-3-319-53168-7_6

10. Newcomb WW, Homa FL, Thomsen DR, et al. Assembly of the herpes simplex virus capsid: characterization of intermediates observed during cell-free capsid formation. J Mol Biol. 1996;263(3):432–446. doi:10.1006/jmbi.1996.0587

11. Heymann JB, Cheng N, Newcomb WW, Trus BL, Brown JC, Steven AC. Dynamics of herpes simplex virus capsid maturation visualized by time-lapse cryo-electron microscopy. Nat Struct Biol. 2003;10(5):334–341. doi:10.1038/nsb922

12. Aksyuk AA, Newcomb WW, Cheng N, et al. Subassemblies and asymmetry in assembly of herpes simplex virus procapsid. mBio. 2015;6(5):e01525–15. Published 2015 Oct 6. doi:10.1128/mBio.01525-15

13. Schrag JD, Prasad BV, Rixon FJ, Chiu W. Three-dimensional structure of the HSV1 nucleocapsid. Cell. 1989;56(4):651–660. doi:10.1016/0092-8674(89)90587-4

14. Booy FP, Newcomb WW, Trus BL, Brown JC, Baker TS, Steven AC. Liquid-crystalline, phage-like packing of encapsidated DNA in herpes simplex virus. Cell. 1991;64(5):1007–1015. doi:10.1016/0092-8674(91)90324-r

15. Gibson W, Roizman B. Proteins specified by herpes simplex virus. 8. Characterization and composition of multiple capsid forms of subtypes 1 and 2. J Virol. 1972;10(5):1044-1052. doi:10.1128/JVI.10.5.1044-1052.1972

16. Gao M, Matusick-Kumar L, Hurlburt W, et al. The protease of herpes simplex virus type 1 is essential for functional capsid formation and viral growth. J Virol. 1994;68(6):3702–3712. doi:10.1128/JVI.68.6.3702-3712.1994

17. Newcomb WW, Trus BL, Cheng N, et al. Isolation of herpes simplex virus procapsids from cells infected with a protease-deficient mutant virus. J Virol. 2000;74(4):1663–1673. doi:10.1128/jvi.74.4.1663-1673.2000

18. White CA, Stow ND, Patel AH, Hughes M, Preston VG. Herpes simplex virus type 1 portal protein UL6 interacts with the putative terminase subunits UL15 and UL28. J Virol. 2003;77(11):6351–6358. doi:10.1128/jvi.77.11.6351-6358.2003

19. Wu W, Newcomb WW, Cheng N, Aksyuk A, Winkler DC, Steven AC. Internal Proteins of the Procapsid and Mature Capsids of Herpes Simplex Virus 1 Mapped by Bubblegram Imaging. J Virol. 2016;90(10):5176–5186. Published 2016 Apr 29. doi:10.1128/JVI.03224-15

20. Tsurumi S, Watanabe T, Iwaisako Y, Suzuki Y, Nakano T, Fujimuro M. Kaposi’s sarcoma-associated herpesvirus ORF17 plays a key role in capsid maturation. Virology. 2021;558:76–85. doi:10.1016/j.virol.2021.02.009

21. Heming JD, Huffman JB, Jones LM, Homa FL. Isolation and characterization of the herpes simplex virus 1 terminase complex. J Virol. 2014;88(1):225–236. doi:10.1128/JVI.02632-13

22. Yang Y, Yang P, Wang N, et al. Architecture of the herpesvirus genome-packaging complex and implications for DNA translocation. Protein Cell. 2020;11(5):339–351. doi:10.1007/s13238-020-00710-0

23. Addison C, Rixon FJ, Preston VG. Herpes simplex virus type 1 UL28 gene product is important for the formation of mature capsids. J Gen Virol. 1990;71 (Pt 10):2377–2384. doi:10.1099/0022-1317-71-10-2377

24. Tengelsen LA, Pederson NE, Shaver PR, Wathen MW, Homa FL. Herpes simplex virus type 1 DNA cleavage and encapsidation require the product of the UL28 gene: isolation and characterization of two UL28 deletion mutants. J Virol. 1993;67(6):3470–3480. doi:10.1128/JVI.67.6.3470-3480.1993

25. Chiu SH, Wu MC, Wu CC, et al. Epstein-Barr virus BALF3 has nuclease activity and mediates mature virion production during the lytic cycle. J Virol. 2014;88(9):4962–4975. doi:10.1128/JVI.00063-14

26. Iwaisako Y, Watanabe T, Hanajiri M, Sekine Y, Fujimuro M. Kaposi’s Sarcoma-Associated Herpesvirus ORF7 Is Essential for Virus Production. Microorganisms. 2021;9(6):1169. Published 2021 May 28. doi:10.3390/microorganisms9061169

27. Gardner MR, Glaunsinger BA. Kaposi’s Sarcoma-Associated Herpesvirus ORF68 Is a DNA Binding Protein Required for Viral Genome Cleavage and Packaging. J Virol. 2018;92(16):e00840–18. Published 2018 Jul 31. doi:10.1128/JVI.00840-18

28. Didychuk AL, Gates SN, Gardner MR, Strong LM, Martin A, Glaunsinger BA. A pentameric protein ring with novel architecture is required for herpesviral packaging. Elife. 2021;10:e62261. Published 2021 Feb 8. doi:10.7554/eLife.62261

29. Draganova EB, Valentin J, Heldwein EE. The Ins and Outs of Herpesviral Capsids: Divergent Structures and Assembly Mechanisms across the Three Subfamilies. Viruses. 2021;13(10):1913. Published 2021 Sep 23. doi:10.3390/v13101913

30. Deng B, O’Connor CM, Kedes DH, Zhou ZH. Cryo-electron tomography of Kaposi’s sarcoma-associated herpesvirus capsids reveals dynamic scaffolding structures essential to capsid assembly and maturation. J Struct Biol. 2008;161(3):419–427. doi:10.1016/j.jsb.2007.10.016

31. Ann M. Arvin, Allison Abendroth. 2021. Varicella-Zoster Virus, p 445–488. Howley, Peter M.; Knipe, David M. (ed), Fields virology, 7th ed, vol 2 Lippincott Williams & Wilkins, Philadelphia, PA.

32. Dai X, Gong D, Lim H, Jih J, Wu TT, Sun R, Zhou ZH. Structure and mutagenesis reveal essential capsid protein interactions for KSHV replication. Nature. 2018 Jan 25;553(7689):521-525. doi: 10.1038/nature25438. Epub 2018 Jan 17. PMID: 29342139; PMCID: PMC6039102.

33. Nealon K, Newcomb WW, Pray TR, Craik CS, Brown JC, Kedes DH. Lytic replication of Kaposi’s sarcoma-associated herpesvirus results in the formation of multiple capsid species: isolation and molecular characterization of A, B, and C capsids from a gammaherpesvirus. J Virol. 2001 Mar;75(6):2866–78. doi: 10.1128/JVI.75.6.2866-2878.2001. PMID: 11222712; PMCID: PMC115913.

34. Goldner T, Hewlett G, Ettischer N, Ruebsamen-Schaeff H, Zimmermann H, Lischka P. The novel anticytomegalovirus compound AIC246 (Letermovir) inhibits human cytomegalovirus replication through a specific antiviral mechanism that involves the viral terminase. J Virol. 2011;85(20):10884–10893. doi:10.1128/JVI.05265-11

35. Dünn-Kittenplon DD, Kalt I, Lellouche JM, Sarid R. The KSHV portal protein ORF43 is essential for the production of infectious viral particles. Virology. 2019;529:205–215. doi:10.1016/j.virol.2019.01.028

36. Perkins EM, Anacker D, Davis A, Sankar V, Ambinder RF, Desai P. Small capsid protein pORF65 is essential for assembly of Kaposi’s sarcoma-associated herpesvirus capsids. J Virol. 2008;82(14):7201–7211. doi:10.1128/JVI.00423-08

37. Naniima P, Naimo E, Koch S, et al. Assembly of infectious Kaposi’s sarcoma-associated herpesvirus progeny requires formation of a pORF19 pentamer. PLoS Biol. 2021;19(11):e3001423. Published 2021 Nov 4. doi:10.1371/journal.pbio.3001423

38. Baines JD, Cunningham C, Nalwanga D, Davison A. The U(L)15 gene of herpes simplex virus type 1 contains within its second exon a novel open reading frame that is translated in frame with the U(L)15 gene product. J Virol. 1997;71(4):2666–2673. doi:10.1128/JVI.71.4.2666-2673.1997

39. al-Kobaisi MF, Rixon FJ, McDougall I, Preston VG. The herpes simplex virus UL33 gene product is required for the assembly of full capsids. Virology. 1991;180(1):380–388. doi:10.1016/0042-6822(91)90043-b

40. Miller JT, Zhao H, Masaoka T, et al. Sensitivity of the C-Terminal Nuclease Domain of Kaposi’s Sarcoma-Associated Herpesvirus ORF29 to Two Classes of Active-Site Ligands. Antimicrob Agents Chemother. 2018;62(10):e00233–18. Published 2018 Sep 24. doi:10.1128/AAC.00233-18

41. Tischer BK, Smith GA, Osterrieder N. En passant mutagenesis: a two step markerless red recombination system. Methods Mol Biol. 2010;634:421–430. doi:10.1007/978-1-60761-652-8_30

42. Brulois KF, Chang H, Lee AS, et al. Construction and manipulation of a new Kaposi’s sarcoma-associated herpesvirus bacterial artificial chromosome clone. J Virol. 2012;86(18):9708–9720. doi:10.1128/JVI.01019-12

43. Watanabe T, Nishimura M, Izumi T, Kuriyama K, Iwaisako Y, Hosokawa K, Takaori-Kondo A, Fujimuro M. Kaposi’s Sarcoma-Associated Herpesvirus ORF66 Is Essential for Late Gene Expression and Virus Production via Interaction with ORF34. J Virol. 2020 Jan 6;94(2):e01300–19. doi: 10.1128/JVI.01300-19. PMID: 31694948; PMCID: PMC6955251.

44. Sugimoto A, Abe Y, Watanabe T, Hosokawa K, Adachi J, Tomonaga T, Iwatani Y, Murata T, Fujimuro M. The FAT10 post-translational modification is involved in the lytic replication of Kaposi’s sarcoma-associated herpesvirus. J Virol. 2021 Feb 24;95(10):e02194–20. doi: 10.1128/JVI.02194-20. Epub ahead of print. PMID: 33627385; PMCID: PMC8139669.

45. Campbell M, Watanabe T, Nakano K, Davis RR, Lyu Y, Tepper CG, Durbin-Johnson B, Fujimuro M, Izumiya Y. KSHV episomes reveal dynamic chromatin loop formation with domain-specific gene regulation. Nat Commun. 2018 Jan 4;9(1):49. doi: 10.1038/s41467-017-02089-9. PMID: 29302027; PMCID: PMC5754359.

46. Fujimuro M, Wu FY, ApRhys C, Kajumbula H, Young DB, Hayward GS, Hayward SD. A novel viral mechanism for dysregulation of beta-catenin in Kaposi’s sarcoma-associated herpesvirus latency. Nat Med. 2003 Mar;9(3):300–6. doi: 10.1038/nm829. Epub 2003 Feb 18. PMID: 12592400.

